# Long non-coding RNA TUG1 is down-regulated in Friedreich’s ataxia

**DOI:** 10.1101/2023.09.22.558879

**Authors:** Mert Koka, Hui Li, Rumana Akther, Susan Perlman, Darice Wong, Brent L Fogel, David R Lynch, Vijayendran Chandran

## Abstract

Friedreich’s Ataxia (FRDA) is a neurodegenerative disorder caused by reduced frataxin (FXN) levels. It leads to motor and sensory impairments and has a median life expectancy of around 35 years. As the most common inherited form of ataxia with no cure, FRDA lacks reliable, non-invasive biomarkers, prolonging and inflating the cost of clinical trials. This study identifies long non-coding RNA Tug1 as a potential blood-based FRDA biomarker.

In a previous study using a frataxin knockdown mouse model (FRDAkd), we observed several hallmark FRDA symptoms and abnormalities in various tissues. Building on this, we hypothesized that a dual-source approach—comparing the data from peripheral blood samples from FRDA patients with tissue samples from affected areas in FRDAkd mice, tissues usually unattainable from patients—would effectively identify robust biomarkers.

A comprehensive reanalysis was conducted on gene expression data from 183 age- and sex-matched peripheral blood samples of FRDA patients, carriers, and controls, as well as 192 tissue datasets from FRDAkd mice. Blood and tissue samples underwent RNA isolation and qRT-PCR, and frataxin knockdown was confirmed through ELISA. Tug1 RNA interaction was explored via RNA pull-down assays. Validation was performed in serum and blood samples on an independent set of 45 healthy controls, 45 FRDA patients; 66 heterozygous carriers, and 72 FRDA patients. Tug1 and Slc40a1 emerged as potential blood-based biomarkers, confirmed in the FRDAkd mouse model (One-way ANOVA, p ≤ 0.05).

Tug1 was consistently downregulated after Fxn knockdown and correlated strongly with Fxn levels (R^2^ = 0.71 during depletion, R^2^ = 0.74 during rescue). Slc40a1 showed a similar but tissue-specific pattern. Further validation of Tug1’s downstream targets strengthened its biomarker candidacy. In additional human samples, TUG1 levels were significantly downregulated in both whole blood and serum of FRDA patients compared to controls (Wilcoxon signed-rank test, p < 0.05). Regression analyses revealed a negative correlation between TUG1 levels and disease onset (p < 0.0037), and positive correlations with disease duration and Functional Disability Stage score (p < 0.04). This suggests that elevated TUG1 levels correlate with earlier onset and more severe cases.

In summary, this study highlights Tug1 as a crucial blood-based biomarker for FRDA. Tug1’s consistent expression variance across human and mouse tissues is closely associated to disease severity and key FRDA pathways. It also correlates strongly with Fxn levels, making it a promising early, non-invasive marker. TUG1 offers potential for FRDA monitoring and therapeutic development, warranting further clinical research.

## Introduction

Friedreich’s ataxia (FRDA) is a debilitating neurodegenerative disorder characterized by the progressive degeneration of the nervous system, resulting in significant motor and sensory impairments^1,2^. The disease ultimately leads to severe physical disability and reduced life expectancy, with the median age of death being 35 years^3,4^. FRDA, being the most common inherited ataxia, is caused by a guanine-adenine-adenine (GAA) trinucleotide repeat expansion in the first intron of the frataxin (FXN) gene^1,2^. This genetic change has a strong correlation with disease severity and its onset^5^. Notably, while heterozygous carriers of the GAA expansion remain asymptomatic, their prevalence ranges from 1:60 to 1:110 in European populations^6,7^. The need for an effective cure for FRDA is challenged by the extensive and expensive clinical trials necessary for drug validation^8^. Therefore, there is an increased emphasis on identifying molecular biomarkers that can quickly monitor disease progression. These biomarkers may expedite evaluations of possible treatments, enhancing patient care and prognosis^8^.

Recent advancements in FRDA research have enabled the assessment of over 20 potential therapeutic interventions in clinical trials^8^. The Food and Drug Administration (FDA) recently approved omaveloxolone, the first drug for FRDA^9^. However, it has shown only moderate efficacy and is associated with some side effects^9^. Nevertheless, a definitive cure or better treatment options for the condition remain elusive. A significant obstacle to the development of effective therapies is the time-consuming and expensive nature of these trials^8^. This underscores the urgency for dependable molecular biomarkers that can expedite evidence-based evaluations of potential treatments. While a simple blood-based assay for FXN could provide benefits like easy sampling, reduced invasiveness, and cost-effectiveness, its creation has been problematic^8^. This is because the active form of FXN is not detectable in serum or plasma; it predominantly resides in erythrocytes^8^. A novel technique utilizing stable isotope dilution Liquid Chromatography-Mass Spectrometry to target the isoform E of FXN from whole blood has been developed^8,10^. However, questions remain regarding its cost-effectiveness, and it is mainly used to assess the efficacy of interventions that aim to increase FXN levels. Therefore, identifying biomarkers that correlate with FXN levels or disease progression is of paramount importance. Discovering such biomarkers in FRDA has been challenging, primarily because of difficulties accessing affected tissues at various disease stages and the lack of animal models that faithfully mimic human disease manifestations.

In our prior research, we presented the frataxin knockdown mouse model (FRDAkd), which closely replicates the symptoms seen in FRDA patients, offering insights into disease progression and the potential for recovery when frataxin expression is restored^11^. In this model, where Fxn expression is reduced, key FRDA symptoms are evident, including ataxia, early mortality, muscle atrophy, degeneration of the dorsal root ganglia, impaired structural integrity of spinal cord axons and myelin, cardiomyopathy, and iron overload^11^. Utilizing this model, this study aims to address the absence of a high confidence molecular biomarker for FRDA. We hypothesize that through the integrative analysis of genomic data sets from (1) FRDA patient samples and (2) samples from the tissues of FRDAkd mice, which are primarily affected in FRDA patients, we can identify robust biomarkers intricately associated with the severity and progression of the disease.

In this study, we focus on the long non-coding RNA (lncRNA) taurine up-regulated gene 1 (TUG1) as a prospective biomarker for FRDA. By analyzing genomics data from both human and mouse models, we detected reduced expression of TUG1 in FRDA. Our results reveal a significant correlation between TUG1 and FXN levels, highlighting the association of TUG1 downregulation with the disease. We suggest that evaluating TUG1 levels could provide a non-invasive metric for disease onset, progression, and severity. Such a biomarker could significantly enrich the therapeutic landscape of FRDA, enabling prompt therapeutic interventions and improving long-term evaluations. In this paper, we report these findings and discuss their implications for the prognosis and management of FRDA. These observations, based on rigorous genomics analysis and a deep understanding of FRDA biology, underscore the potential of lncRNA TUG1 as a promising molecular biomarker in FRDA.

## Materials and methods

### Analysis of Human and Mouse Gene Expression Data

A total of 733 peripheral blood samples from FRDA patients (411), carriers (228), and controls (94) were reanalyzed to identify blood-based biomarker for FRDA (GEO dataset: GSE102008)^12^. To avoid any confounding effect in this data series, we conducted differential gene expression analyses on age- and sex-matched 183 samples consisting of 72 patients, 68 carrier, and 43 controls (**Figure 1A**). For the mouse data, a total of 192 microarray datasets from 64 RNA samples derived from FRDAkd mice were analyzed to compare and examine the overlap of the differentially expressed genes obtained from FRDA patient data and mouse data (GEO dataset: GSE98790)^11^. Both raw data were log transformed and checked for outliers. Inter-array Pearson correlation and clustering based on variance were used as quality-control measures. Quantile normalization was used and contrast analysis of differential expression was performed by using the LIMMA package (RRID:SCR_010943). Briefly, a linear model was fitted across the dataset, contrasts of interest were extracted, and differentially expressed genes for each contrast were selected using an empirical Bayes test statistic^13^.

**Figure 1.**
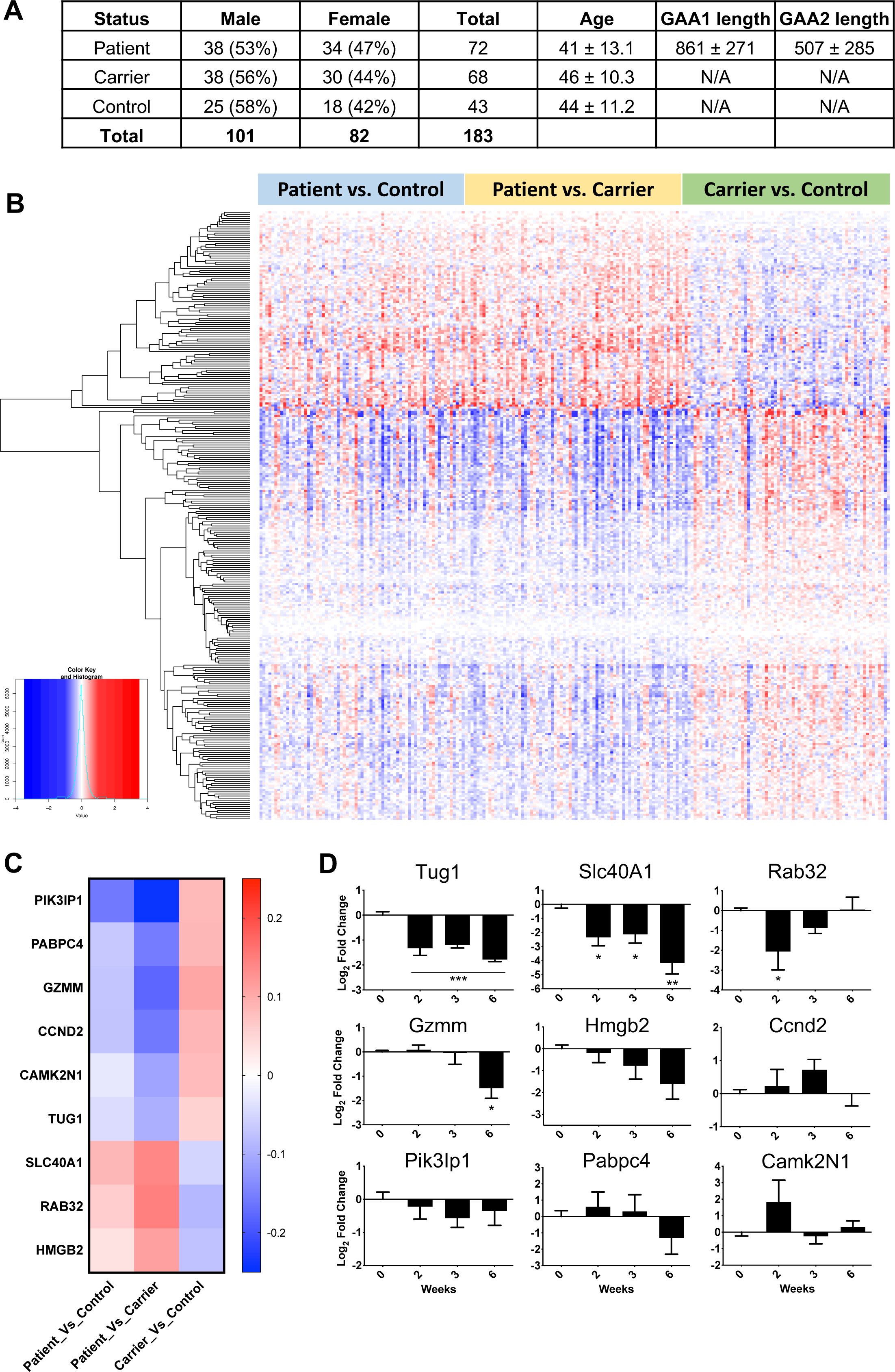
Differential Gene Expression in Whole Blood of FRDA Patients. (**A**) Demographic breakdown of the participants, featuring sex and age-matched samples used for the analysis. (**B**) A heat map representing significant gene up- and down-regulation (depicted in rows) in whole blood from FRDA patients (n = 72), carriers (n = 68), and controls (n = 43), sourced from GEO dataset GSE102008. Red and blue color intensities correspond to gene up- and down-regulation, respectively. (**C**) A heat map showing the top nine differentially expressed genes in FRDA patient’s whole blood, selected for subsequent validation. (**D**) Preliminary screening for expression levels of candidate genes in the whole blood of FRDA-knockdown mice. Expression levels of candidate genes, normalized to Hprt1, in the whole blood of FRDA-knockdown mice post-treatment with doxycycline from week 0 to week 6, were examined using relative quantitative PCR analyses. Up- and down-regulation is indicated by positive and negative signals, respectively. N = 3-5. One-way ANOVA and Welch’s t-test were employed for statistical analysis. Data are presented as mean ± SEM, with significance marked as * = p < 0.05, ** = p < 0.01, *** = p < 0.001.

### Animal Study Design and Ethics

All animal experiments were carried out in accordance with relevant guidelines and regulations, and upon the approval of an Institutional Animal Care and Use Committee (IACUC) at the University of Florida (protocol number # 201909663) and in compliance with the ARRIVE guidelines. Wildtype mice were C57BL/6J from the Jackson Lab, and transgenic mice were FRDAkd mice in C57BL/6J background. Mice were sorted into four different groups: wildtype with doxycycline (Wt + dox), transgenic with doxycycline (Tg + dox), without doxycycline (Tg – dox) and transgenic with dox removal - rescue (Tg + res). Number of mice (N) was 4 for each time point. Male to female ratio was 1:1 for all groups. Animals were to be euthanized at weeks 0, 2, 3, 4, 6, 8, R1, R2, R3, R4, R6 and R8 (R= Rescue – Dox removal). Transgenic with dox and wildtype with dox animals were given 2000 mg/kg doxycycline hyclate (∼87% doxycycline) diet, TD.09633 from Envigo. Transgenic without dox animals were given normal diet. Transgenic with rescue were switched from doxycycline hyclate diet to normal diet.

### Mice Tissue and Blood Sample Collection Protocol

Animals were deeply anesthetized with isoflurane, and cardiac puncture was performed to collect the blood in EDTA covered RNAse-free tube. The blood was then centrifuged at 3000 g for 15 minutes. Supernatant (the plasma) was collected and snap frozen with liquid nitrogen. Mice were placed on petri dish on ice and dissection of heart, liver, brain, muscle from femur were performed. Spinal column was isolated, and hydraulic extrusion of spinal cord was performed with ice cold 1X PBS. All tissues were cut and rinsed in 1X cold PBS on ice and quickly transferred over to 1.7 mL sterile, RNAse-free tubes to be snap frozen with liquid nitrogen. All samples were stored in -80°C.

### RNA Isolation, cDNA Preparation, and qRT-PCR Protocols

Tissue samples of blood, muscle, heart, spinal cord, brain, and liver from FRDAkd mice were retrieved from a -80°C freezer for RNA isolation. The PAXgene blood RNA kit (cat #762164) was used for blood samples, while the miRNeasy mini kit (cat #2170040) was utilized for other tissues. SuperScript VILO master mix was employed to synthesize cDNA from the extracted RNA. Depending on the tissue type, total RNA concentrations ranging from 0.5-1.0 µg were used to prepare 10 µL cDNA aliquots as per the manufacturer’s guidelines. For qRT-PCR, iTaq Universal SYBR Green Supermix was used. Each well contained 0.2 µL of the cDNA prep and 500 nM each of forward and reverse primers. The qRT-PCR was conducted using the Bio-Rad CFX96 real-time PCR system. Melt curve analysis was performed to confirm the presence of a single amplicon for each of the following primer sets.

#### List of primers

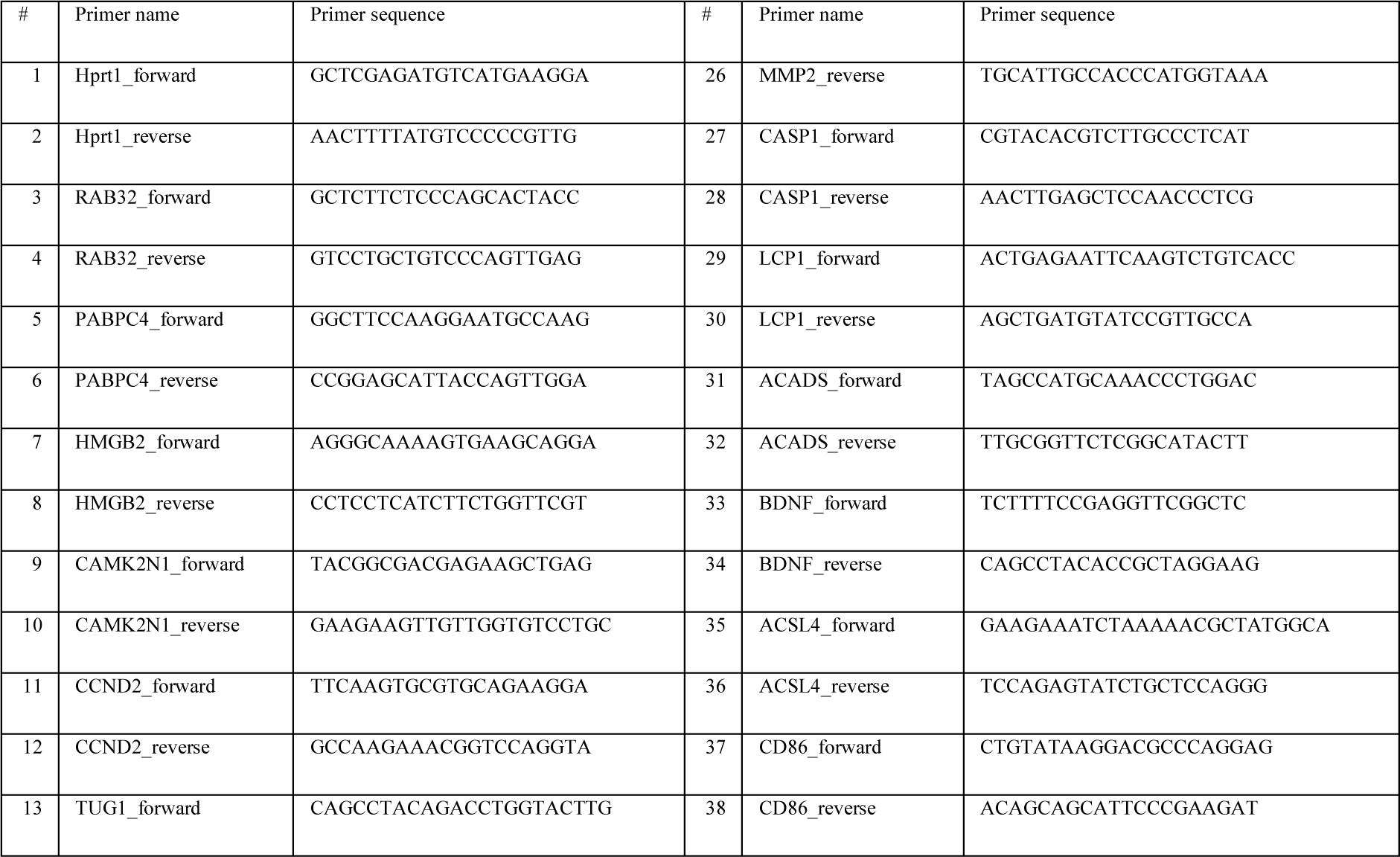
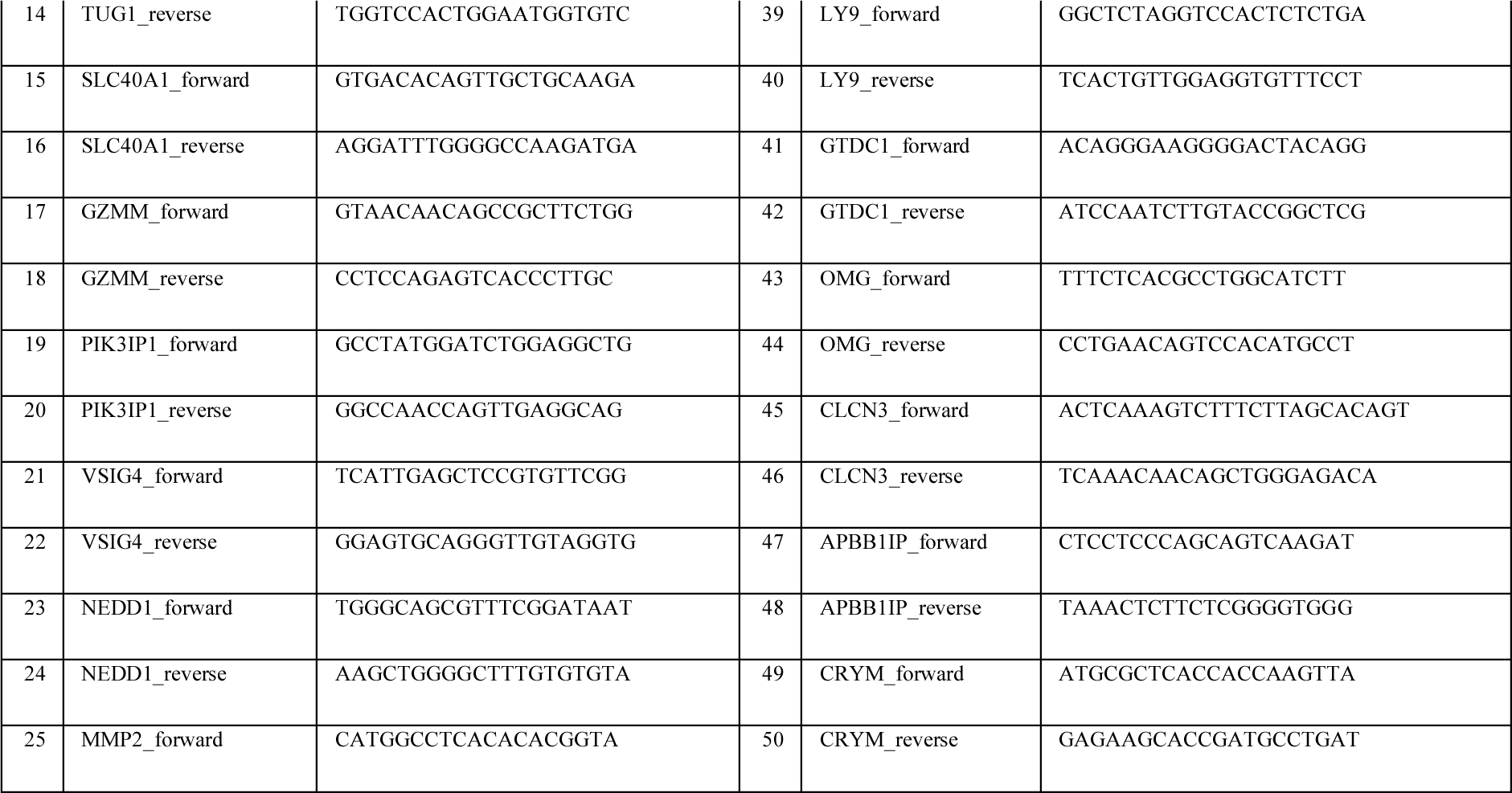

### Validation of Frataxin Knockdown Using ELISA in Heart Tissue

To validate the knockdown of frataxin, heart tissue samples were selected for ELISA analysis. Snap-frozen heart tissue aliquots from all 40 mice were retrieved from -80°C storage and weighed in new tubes. For protein extraction, 10 µL of 1X ELISA lysis buffer (Abcam ab176112 ELISA kit) per mg of tissue was used in conjunction with a pestle tissue homogenizer. Protein concentration was quantified using the BCA assay kit (cat #23225). Subsequently, 30 µg of total protein from each heart sample was loaded into ELISA wells, and the protocol outlined in the ELISA kit was followed. Frataxin concentrations in the samples were interpolated using a second-order polynomial regression model based on a standard curve generated from human lyophilized recombinant frataxin protein provided in the kit. Data were plotted using Prism 6.

### RNA Pull-Down Assay to Investigate Tug1 RNA Interactome

To explore the RNA interactome of Tug1, a previously established RNA pull-down protocol for long non-coding RNAs (lncRNAs) was employed^14^. The secondary structure of lncRNA Tug1 was predicted using RNAstructure Webserver, and an antisense biotinylated DNA oligonucleotide probe with minimal probability of internal base pairing was designed via the Fold algorithm^15^. A non-specific DNA oligonucleotide probe served as a negative control. Heart tissues were initially crosslinked with paraformaldehyde, lysed and sonicated. Lysates were mixed with hybridization buffer, and an input sample was frozen until use. Subsequent hybridization involved adding biotinylated DNA probes to the lysates for 4 hours, followed by overnight incubation with streptavidin beads. The beads were magnetically separated and washed several times. Proteinase K treatment was performed before RNA isolation using the miRNeasy mini kit (cat #2170040). Reverse transcription was carried out with SuperScript IV VILO master mix with ezDNase (cat #11766050). The assay was analyzed using RT-qPCR to determine relative enrichment of target molecules in comparison to input samples.

### Clinical Sample Acquisition and Ethical Approvals

Acquisition protocols for human clinical samples were approved by the Institutional Review Boards (IRBs) at the University of California, Los Angeles (UCLA) and the Children’s Hospital of Philadelphia (CHOP). Informed written consent was obtained from all participants, or from legal guardians in the case of subjects under 18 years. The sample set comprised 45 healthy controls and 45 FRDA patients for serum samples. Additionally, blood samples were collected from 66 heterozygous carriers of the FXN gene with GAA-repeats (unaffected) and 72 FRDA patients.

### Venous Blood and Serum Sample Collection and Storage

Blood samples were collected from each participant using 8.5 mL purple-cap Vacutainer EDTA tubes (Becton, Dickinson and Company, NJ, USA, Catalog #367861). The tubes were gently inverted 8–10 times post-collection for anticoagulant mixing. The samples were transferred into 2.4 mL Eppendorf tubes (Eppendorf AG, Hamburg, Germany) with sterile pipettes and immediately stored at -80°C in an ultra-low freezer. For serum, blood samples were allowed to clot at room temperature for 30 minutes before undergoing centrifugation at 1500 × g for 10 minutes at 4°C using a refrigerated centrifuge (Eppendorf AG, Hamburg, Germany). The supernatant serum was pipetted into sterile 1.5 mL Eppendorf tubes and stored at -80°C until further analysis.

### Statistical Analysis and Data Interpretation

Data normality was assessed using Kolmogorov–Smirnov, Shapiro–Wilk, or D’Agostino-Pearson tests based on the dataset size. For normally distributed data, one-way or two-way ANOVA was employed for statistical analysis. In cases where the data did not conform to a normal distribution, alternative statistical tests were applied as indicated in the respective figure legends. Grouped data were analyzed using one or two-way ANOVA with appropriate post hoc multiple comparison tests. All statistical analyses were conducted using GraphPad Prism 9 software (GraphPad Prism, RRID: SCR_002798). A p-value of ≤ 0.05 was considered statistically significant. Results were calculated based on N = 4 or more per time point and are presented as mean ± SEM.

### Data availability

The raw gene expression data for FRDA patients can be accessed via the NCBI Gene Expression Omnibus (GEO) database, under the accession number GSE102008. Similarly, gene expression datasets for FRDAkd mice are publicly available on the GEO database under the accession number GSE98790. All additional data pertinent to this study can be obtained upon reasonable request directed to the corresponding author.

## Results

### Blood-based Biomarker Discovery in FRDA

In this study, we reanalyzed an extensive series of previously published unbiased gene expression profiles comprising 733 individuals, including 411 FRDA patients, 228 carriers, and 94 controls, with the aim of identifying blood-based biomarkers specific to FRDA^12^. Differential gene expression analyses were conducted on age- and sex-matched 183 samples, made up of 72 patients, 68 carriers, and 43 controls, to eliminate any confounding effects in these data series (**Figure 1A**). This analysis identified 293 genes that were differentially expressed with an FDR of less than 10% (**Figure 1B**). Subsequent functional annotation analysis of these transcripts revealed predominant Gene Ontology categories, such as adaptive and innate immune response. This finding aligns with previous research, where immune system activation was one of the earliest regulated pathways post Fxn knockdown^11,12^. Thus, utilizing the differentially expressed 293-gene list derived from FRDA patient whole blood data, we postulate that it is feasible to rank and validate these genes for the identification of potential biomarkers for FRDA.

### Biomarker Validation of Tug1 and Slc40a1 in FRDA Knockdown Mice

Utilizing gene expression data from the FRDAkd mouse model^11^, we examined the overlap of the 293 differentially expressed genes obtained from FRDA patient data and mouse data. This analysis focused on three tissues primarily affected in FRDA: the heart, dorsal root ganglia (DRG) neurons, and the cerebellum. We investigated these tissues after Fxn knockdown, achieved through doxycycline (dox) treatment, and subsequent rescue following dox removal^11^. We found 49 genes that were differentially expressed in both FRDA patient data and FRDAkd mouse gene expression data, with an FDR of less than 5%. These genes were then ranked based on consistency across samples and a non-parametric Kruskal-Wallis p-value less than 1.2 x 10^-4^, and the top 9 genes (Slc40A1, Rab32, Tug1, Pabpc4, Gzmm, Hmgb2, Camk2N1, Ccnd2, and Pik3Ip1) were selected for further validation (**Figure 1C**). To validate these 9 genes and determine if these candidate biomarkers are direct targets due to Fxn knockdown, we performed a focused examination. Specifically, we assessed whether they were differentially expressed in the whole blood as early as two weeks after Fxn knockdown in FRDAkd mice. This process aimed to discern if the observed alterations in gene expression were a direct consequence of changes in Fxn levels, thus strengthening the hypothesis that these genes could serve as potential biomarkers for FRDA. Their expression levels were assessed by qRT-PCR at various intervals (0, 2, 3, and 6 weeks) post dox treatment. Among the 9 genes, taurine-upregulated gene 1 (Tug1) and ferroportin-1 (Slc40a1) were found to be significantly differentially expressed (One-way ANOVA, p ≤ 0.05) as early as two weeks after Fxn knockdown in whole blood (**Figure 1D**). The differential expression of Tug1 and Slc40a1 in peripheral blood positions these genes as exemplary candidate biomarkers for FRDA, particularly appealing due to the minimal invasiveness required for patient sample collection.

### Assessment of Frataxin Levels and Correlation with Candidate Biomarkers in FRDAkd Mouse Model

Carriers and FRDA patients exhibit reduced frataxin levels when compared with controls. The lateral flow immunoassays conducted on buccal cells revealed frataxin protein levels at 50.5% in carriers and 21.1% in FRDA patients^16^. The FRDAkd mouse model facilitates an exploration of various frataxin level profiles across different tissues, mirroring those observed in controls, carriers, and FRDA patients. These profiles vary based on the dose, duration, and rescue of dox treatment. This model afforded the opportunity to investigate the correlation between the expressions of the top two candidate biomarkers (Tug1 and Slc40a1) at different time-points and the varying levels of FXN. Utilizing enzyme-linked immunosorbent assays (ELISA) on heart samples, we demonstrated the FXN level profiles within the FRDAkd mouse model. After dox treatment, the FXN levels in the heart samples of FRDAkd mice were observed to decrease by 47% by week 2, and 93% by week 6 (**Supplementary Figure 1A**). Extended exposure to dox treatment yielded lower FXN levels in the heart, while Tg – Dox and Wt + Dox remained unchanged. At the mRNA level, a significant knockdown of Fxn was detected at week 6 in the heart, muscle, spinal cord, brain, and liver, with evidence of partial rescue of Fxn expression by 8 weeks post-dox removal (R8) in the heart, muscle, brain, and liver (**Figure 2A**).

**Figure 2.**
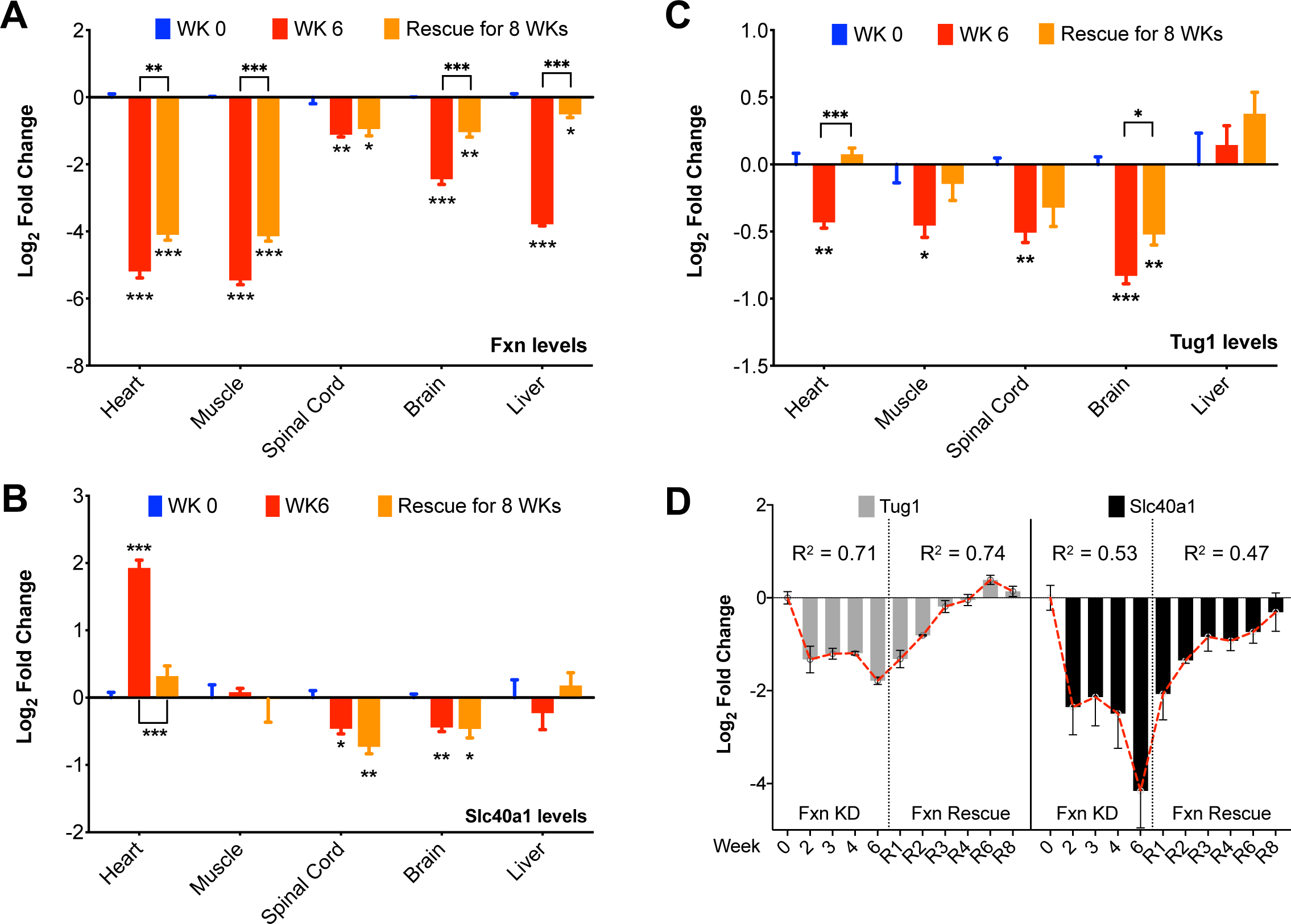
Tissue-Specific Expression Levels of Fxn, Tug1, and Slc40a1 in FRDA-Knockdown Mice. (**A**) Fxn, (**B**) Slc40a1 and (**C**) Tug1, expression levels in heart, muscle, spinal cord, brain, and liver tissues from FRDA-knockdown mice treated with doxycycline for 0 and 6 weeks (Fxn knockdown), followed by an 8-week rescue period (Fxn recovery). Each time point included four samples (N = 4). (**D**) Linear regression models were employed to calculate R^2^ values based on the log_2_ fold change of Tug1 and Slc40a1 expression levels, normalized to Hprt1, in the blood of FRDA-knockdown mice during the Fxn knockdown and rescue phases. N = 3-5. Welch’s t-test was used for statistical analysis. Data are presented as mean ± SEM, with significance marked as * = p < 0.05, ** = p < 0.01, *** = p < 0.001.

We next delved into the examination of Tug1 and Slc40a1 gene expression across various tissues in the FRDAkd mouse model to explore the consistency of expression changes across different tissues. Specifically, we analyzed the gene expression levels of Tug1 and Slc40a1 in five diverse tissues (heart, muscle, spinal cord, brain, and liver) at three time-points: immediately after Fxn knockdown at 0 and 6 weeks post dox treatment, followed by rescue at 8 weeks post dox removal (**Figure 2B and 2C**). We observed, Slc40a1 demonstrated a significant downregulation in spinal cord and brain tissues after Fxn knockdown, and interestingly, the expression of Slc40a1 mRNA was not fully reversed during the Fxn restoration phase, indicating a slow reversal process. Conversely, in the heart tissue, Fxn knockdown led to an upregulation of Slc40a1, with subsequent Fxn rescue resulting in a reversal of this expression pattern. In the liver and muscle tissues, there were no evident alterations in Slc40a1 expression attributed to Fxn knockdown (**Figure 2B**). It is essential to note that ferroportin-1, is a critical transmembrane protein involved in iron export^17^. Previous findings have highlighted iron metabolism dysregulation in heart autopsies of FRDA patients^18,19^. Moreover, iron concentrations in plasma samples were found to be significantly lower in FRDA patients compared to healthy controls^20^. These observations align cohesively with our results, further emphasizing the potential role of Slc40a1 in the pathological mechanisms underlying FRDA.

### Analysis of the Potential of Tug1 as a Specific Biomarker for FRDA

Our next candidate, long non-coding RNA (lncRNA) Tug1, is known to interact with the polycomb repressor complex and functions in the epigenetic regulation of transcription. Tug1 has been shown as a regulatory factor, involved in cellular processes such as cell proliferation^21–23^, apoptosis^24–28^, cell cycle^22,24,29,30^, and mitochondrial bioenergetics^31^. Notably, these processes are also primarily impacted in FRDA. Through our analysis, we discovered that Tug1 was markedly downregulated in all examined tissues following Fxn knockdown, with the exception of the liver. During the Fxn restoration phase, the expression level of Tug1 exhibited complete recovery in the heart and muscle tissues. However, there was only a limited rescue of Tug1 expression in the spinal cord and brain, as depicted in **Figure 2C**. This limited rescue can be attributed to residual levels of dox remaining in the CNS after its withdrawal. In summary, these experiments reveal a consistent expression pattern of Tug1 in response to Fxn knockdown and restoration, highlighting its significant downregulation across various tissues and its potential role as a specific biomarker for FRDA.

### Analysis of Tug1 and Slc40a1 Expression in Whole Blood During Fxn Knockdown and Rescue in FRDAkd Mice

Following the preliminary screening of candidate genes at early time points, we evaluated the expression of Tug1 and Slc40a1 genes in whole blood across 11 specific time points, consisting of 5 stages during Fxn knockdown and 6 stages during Fxn rescue. During the Fxn depletion phase, both Tug1 (R^2^ = 0.71) and Slc40a1 (R^2^ = 0.53) exhibited a strong linear decline. Conversely, during the Fxn rescue phase, Tug1 (R^2^ = 0.74) and Slc40a1 (R^2^ = 0.47) demonstrated a marked linear increase (**Figure 2D**). To control for the potential confounding effects of dox treatment, we examined the expression of Tug1 and Slc40a1 in wildtype mice across 0, 6, and R8 weeks of dox treatment and subsequent removal (**Supplementary Figure 1B**). No significant alterations in mRNA expression of Tug1 and Slc40a1 were observed in wildtype animals during either dox treatment or withdrawal, supporting the notion that the changes in transgenic mice were attributable to changes in Fxn levels. In the context of these findings, Tug1 emerged as a more suitable peripheral biomarker for FRDA compared to Slc40a1.

Our previous gene expression data in FRDAkd mice revealed significant down-regulation of Tug1 in the cerebellum and dorsal root ganglion throughout disease progression, paralleling the trend in Fxn expression (**Supplementary Figure 2A**). In FRDA, significant neuronal loss in the dentate nuclei and extensive cerebellar damage have been reported^4,32,33^. Interestingly, we observed that TUG1 expression is highest in the cerebellum among all human brain regions (**Supplementary Figure 2B**). Furthermore, in FRDAkd mice treated with dox (Fxn knockdown), we observed a downregulation of Tug1 as early as the third week of treatment (**Supplementary Figure 2C**). In summary, Tug1’s strong linear correlation with Fxn levels during knockdown and rescue phases, consistency in expression across various tissues, and early detection of down-regulation in specific regions like the cerebellum present it as a suitable and compelling candidate biomarker for FRDA.

### Tug1 Downstream Targets are Altered in FRDAkd Mice

Tug1 functions as a regulatory factor, known for controlling intricate cellular processes that are also affected in FRDA, including cell proliferation^21–23^, apoptosis^24–28^, cell cycle^22,24,29,30^, and mitochondrial bioenergetics^31^. To further examine the role of Tug1 in FRDA and validate its potential to serve as a biomarker, we investigated the downstream target genes of Tug1. One study identified 630 genes as Tug1 targets using a modified RNA pull-down assay with promoter-microarray analysis in BrU-labelled Tug1-transfected glioma cells^34^. Cross-comparison of these 630 targets of Tug1 combined with 33 experimentally verified Tug1 targets against the list of differentially expressed genes in heart, cerebellum and dorsal root ganglions of FRDAkd mice led to the identification of 40 overlapping genes (**Figure 3A**). To validate this, we manually selected the top sixteen Tug1 targets based on differential expression p-value, expression in multiple tissues, and association with cellular processes affected in FRDA as evidenced by the literature. The selected genes for validation were: Vsig4, Nedd1, Mmp2, Casp1, Lcp1, Acads, Bdnf, Acsl4, Cd86, Ly9, Gtdc1, Omg, Clcn3, Apbb1ip, Crym, and Ccnd2.

**Figure 3.**
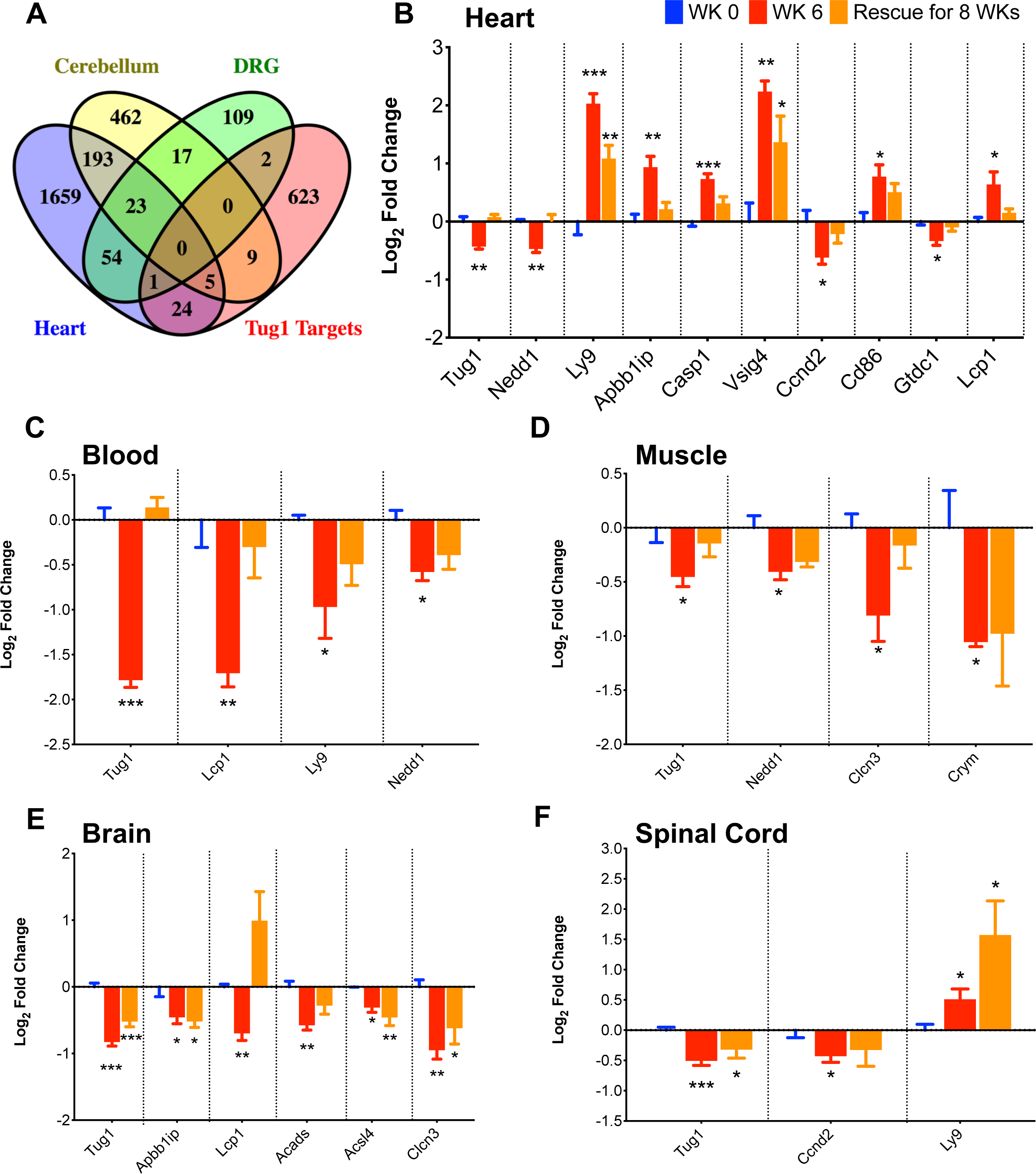
Impact of Tug1 Downregulation on its Target Genes in Various Tissues of FRDA-Knockdown Mice. (**A**) Venn diagram illustrating the cross-comparison of Tug1 targets identified from RNA-pulldown assay using Tug1-transfected human glioma cells and other experimental data, with the list of differentially expressed genes uncovered via transcriptomic analysis in the heart, cerebellum, and dorsal root ganglia (DRG) of FRDA-knockdown mice. (**B - F**) Expression changes in Tug1 targets within (**B**) heart, (**C**) blood, (**D**) muscle, (**E**) brain, and (**F**) spinal cord tissues of FRDA-knockdown mice treated with doxycycline for 0 and 6 weeks (Fxn knockdown), followed by an 8-week rescue phase (Fxn recovery). These alterations were assessed through quantitative reverse transcription PCR experiments. Four samples were taken at each time point (N = 4). One-way ANOVA was used for statistical analysis. Data are presented as mean ± SEM, with significance denoted as * = p < 0.05, ** = p < 0.01, *** = p < 0.001.

We conducted qRT-PCR experiments on blood, heart, muscle, brain, and spinal cord tissues after Fxn knockdown and rescue in FRDAkd mice to validate Tug1 targets. Out of 16 genes tested, 9 in heart, 5 in brain, 3 in muscle, 3 in blood and 2 in spinal cord were differentially expressed. Nedd1, for instance, was significantly downregulated in blood, heart, and muscle in FRDAkd mice treated with dox for six weeks (**Figure 3**). Depletion of Nedd1, a centrosome-localized protein, is linked to senescence induction in mouse embryonic fibroblasts^35^, a phenomenon also observed in frataxin-deficient human neuroblastoma cells^36^. Ccnd2, a protein responsible for regulating cell cycle, was regulated by lncRNA Tug1 in bladder cancer, and the authors also reported that expression of Ccnd2 positively correlated with Tug1 expression^37^. Ccnd2 expression was downregulated in spinal cord and heart. Clcn3, a protein thought to affect vesicle trafficking and exocytosis, was downregulated in muscle and brain. Disrupted Clcn3 leads to impairment of acidification of synaptic vesicles, which causes severe neurodegeneration^38^. With frataxin knockdown and the corresponding Tug1 deficiency, Lcp1 was notably downregulated in blood and brain but upregulated in the heart. Ly9 exhibited downregulation in blood and upregulation in spinal cord and heart. Both Lcp1 and Ly9 have been implicated in T-cell activation^39,40^, essential for immune system regulation. In FRDAkd mice, immune system activation was among the earliest pathways affected after frataxin knockdown^11^. These findings strengthen the case for Tug1 as a promising biomarker for FRDA, given the significant alterations in its downstream targets caused by Fxn knockdown in FRDAkd mice.

### RNA Pulldown Analysis of Tug1 Target Gene Associations in FRDA-Knockdown Heart Tissue

To further elucidate the interaction between Tug1 and its target genes in the heart tissue of FRDA-knockdown mice, we conducted an RNA pulldown experiment. Utilizing a data-driven method with biotinylated probes, we aimed to isolate and specifically identify the target genes binding to Tug1 (**Figure 4A**). A subsequent analysis using quantitative PCR on the Tug1 RNA pulldown samples obtained from the heart tissue of FRDA-knockdown mice shed light on the enrichment of specific genes, reinforcing the interactions between Tug1 and its target genes. Through this meticulous approach, we successfully validated the significant association of Tug1 with several target genes, namely Nedd1, Ccnd2, and Lcp1 (**Figure 4B**). These results not only confirm Tug1’s pivotal role in the molecular mechanisms of FRDA but also underscore its potential as a key factor in therapeutic targeting of FRDA’s pathogenesis.

**Figure 4.**
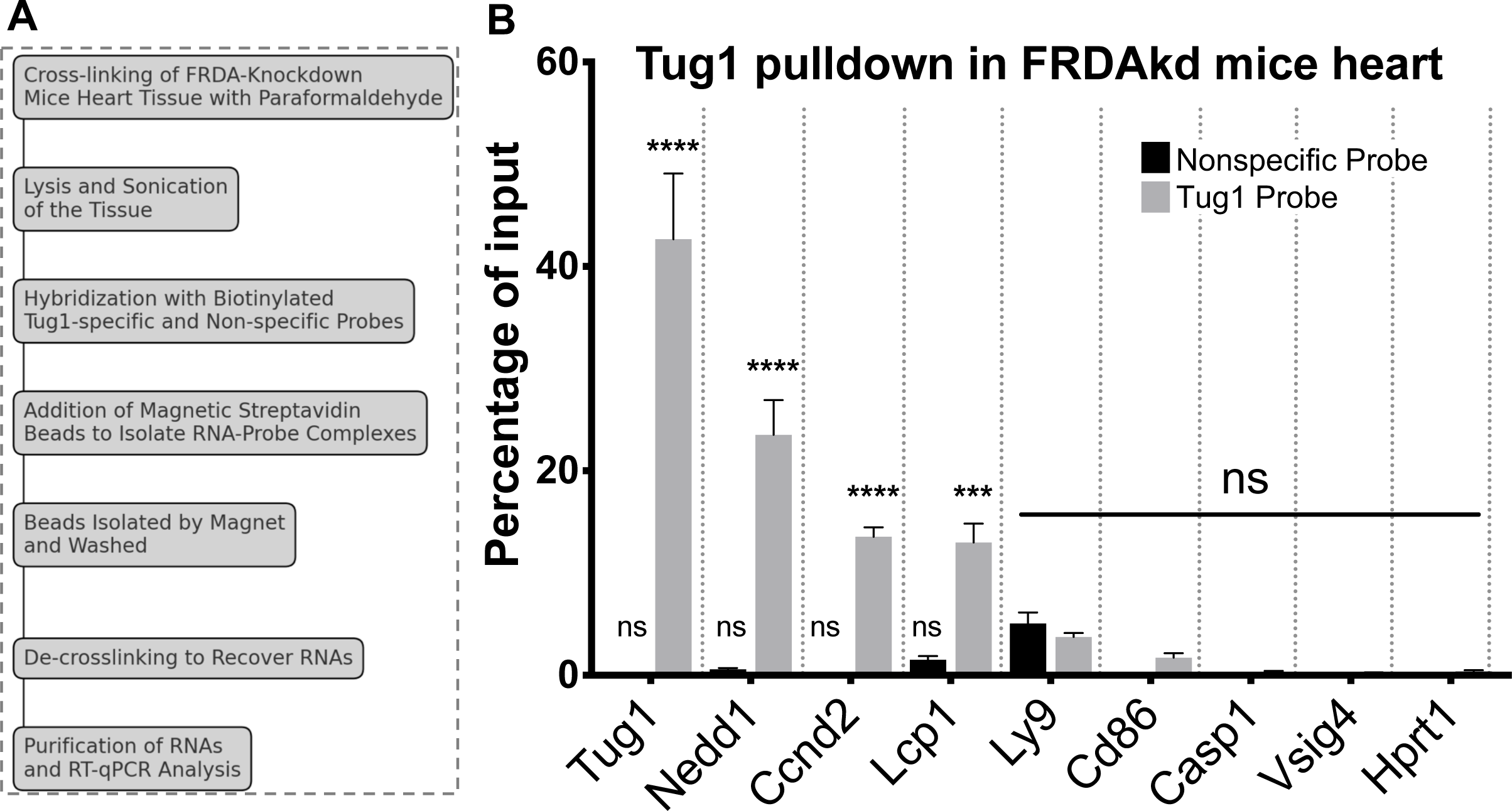
Tug1 Target Enrichment in the Heart Tissue of FRDA-Knockdown Mice Revealed Through RNA Pulldown Experiment. (**A**) A schematic of the RNA pulldown experiment, where biotinylated probes were designed and used to isolate Tug1 and its target genes using streptavidin beads. A non-specific probe served as a negative control. (**B**) Bar graph representing the percentage of input enrichment for several target genes, showing their significant association with Tug1. Each target gene is presented as a separate bar in the graph. These gene enrichments were determined by performing quantitative PCR on the Tug1 RNA pulldown samples derived from the heart tissue of FRDA-knockdown mice. A two-way ANOVA was used to evaluate statistical significance. Data are shown as mean ± SEM, with * = p < 0.05, ** = p < 0.01, *** = p < 0.001 indicating levels of significance.

### Quantification of TUG1 Levels in FRDA Patients and Correlation Analysis with Key Clinical Parameters

To investigate TUG1 expression levels in patients with FRDA, we analyzed the publicly available microarray dataset GSE102008. Our comparative evaluation focused on whole-blood TUG1 expression levels across age- and sex-matched individuals, including FRDA patients (n=72), heterozygous carriers (n=68), and healthy controls (n=43). By employing a one-way ANOVA with Holm-Sidak’s multiple comparisons test, we identified a significant downregulation of TUG1 expression in the FRDA cohort (**Figure 5A**). Given the previous detection of TUG1 from human serum samples in patients with multiple myeloma^41^, we explored its expression in FRDA serum samples. Our analysis of serum samples from 45 patient serum and 45 healthy control serum samples showed a significant downregulation of serum TUG1 expression levels in FRDA patients compared to healthy controls, as confirmed through reverse transcription-quantitative PCR, employing the Wilcoxon signed-rank test (**Figure 5B**). We also examined and quantified the whole-blood TUG1 expression levels between age- and sex-matched FRDA patients (n=72) and heterozygous carriers (n=66). Through reverse transcription-quantitative PCR, we detected a significant downregulation (P<0.05) in comparison to the control group (Wilcoxon signed-rank test) (**Figure 5C**). The results collectively demonstrated a marked downregulation of TUG1 expression in both whole blood and serum from FRDA patients (**Fig. 5A-C**). Subsequently, to uncover the functional relevance of TUG1 expression, we constructed a correlation heatmap that revealed the relationship between blood TUG1 expression levels and various demographic and clinical characteristics of FRDA patients. The correlation studies determined a substantial inverse correlation between TUG1 expression levels and disease onset (**Fig. 5D**). Notably, we detected positive correlations with disease duration and the Functional Disability Stage (FDS) score, clinical parameters known to directly influence each other (**Fig. 5D**). In summary, our thorough examination of TUG1 expression levels across multiple cohorts and experimental frameworks has unveiled a distinct gene expression profile in FRDA patients. This novel understanding not only underscores the potential of TUG1 as a therapeutic target but also lays the groundwork for subsequent investigations into the molecular pathways modulated by TUG1 in FRDA.

**Figure 5.**
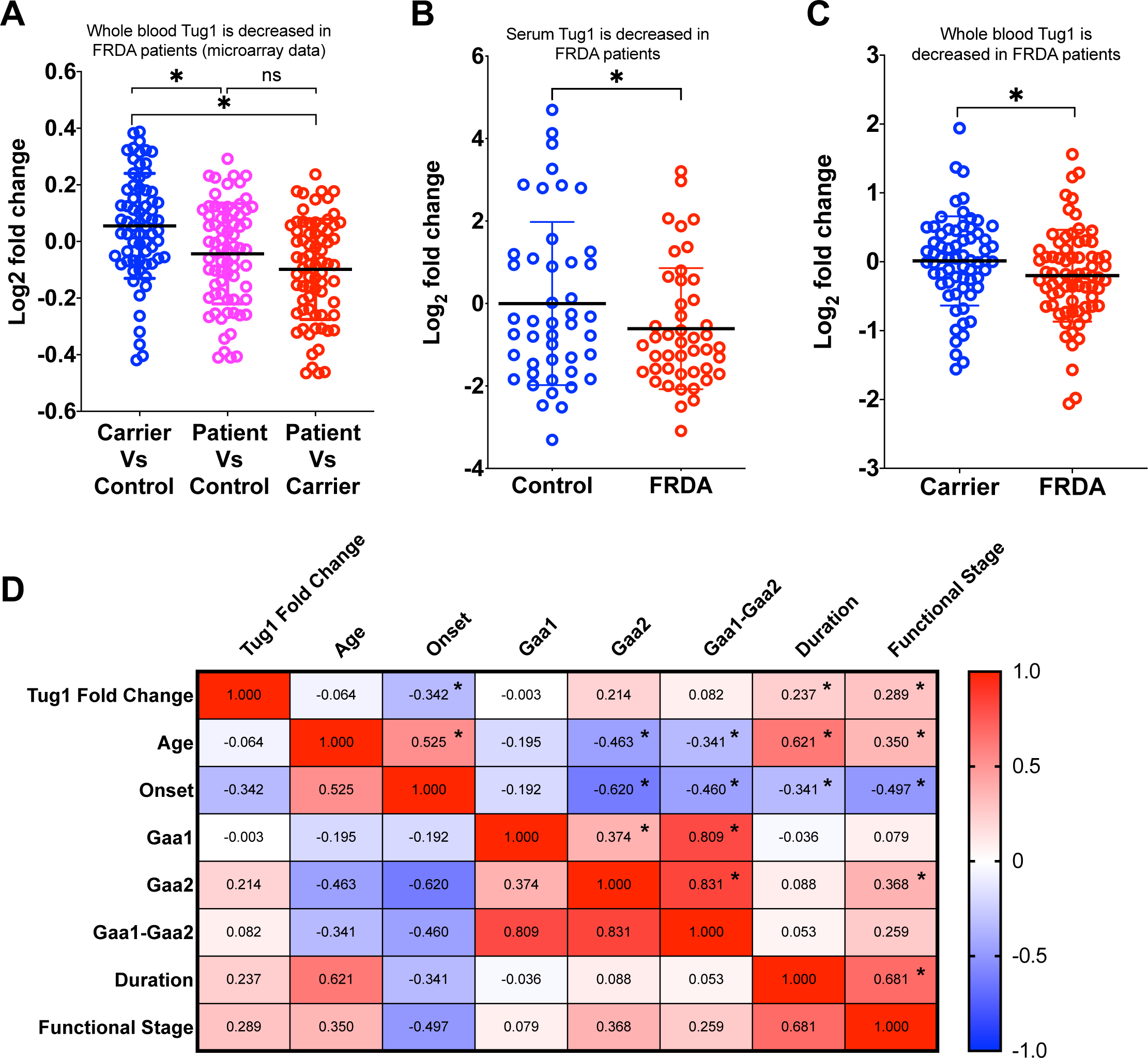
TUG1 Expression Levels are Reduced in FRDA Patients. (**A**) Scatter plot comparing whole-blood TUG1 expression levels among age- and sex-matched FRDA patients (n=72), heterozygous carriers (n=68), and healthy controls (n=43), using the GEO microarray dataset GSE102008. Significance was assessed using one-way ANOVA with Holm-Sidak’s multiple comparisons test. Data points are represented individually, with mean ± SD. * P<0.05. (**B**) Serum TUG1 expression levels in FRDA patients (n=45) exhibit significant downregulation compared to healthy controls (n=45), as determined by reverse transcription-quantitative PCR. Wilcoxon signed-rank test was performed. (**C**) Scatter plot illustrating the comparison of whole-blood TUG1 expression levels between age- and sex-matched FRDA patients (n=72) and heterozygous carriers (n=66) via reverse transcription-quantitative PCR. * P<0.05 vs. control group (Wilcoxon signed-rank test was performed). (**D**) Heatmap displaying the correlation matrix between blood TUG1 expression levels and various demographic characteristics of FRDA patients. Cell coloration ranges from red (indicating positive correlations) to blue (representing negative correlations), reflecting the strength of associations. Pair-wise Pearson correlation coefficients are displayed in each cell, with stars marking significance at p-value < 0.05.

### Regression analyses on TUG1 Expression Levels and FRDA Clinical Variables

Given the evidence indicating significant correlations between TUG1 and various factors including disease onset, duration, and the FDS score in FRDA, we further explored these relationships through linear regression analyses. As expected, a key observation emerging from these analyses was a marked association between the functional disability stage and age, emphasizing the clinical significance of these parameters in predicting the progression of the disease (**Fig. 6A**). Further, the linear regression results validated a negative correlation between Tug1 levels and disease onset (p < 0.0037) (**Fig. 6B**), reinforcing the relevance of Tug1 in understanding disease onset. Additionally, positive correlations were established between Tug1 expression and both disease duration (p < 0.04) and the FDS score (p < 0.04) (**Fig. 6 C, D**).

**Figure 6.**
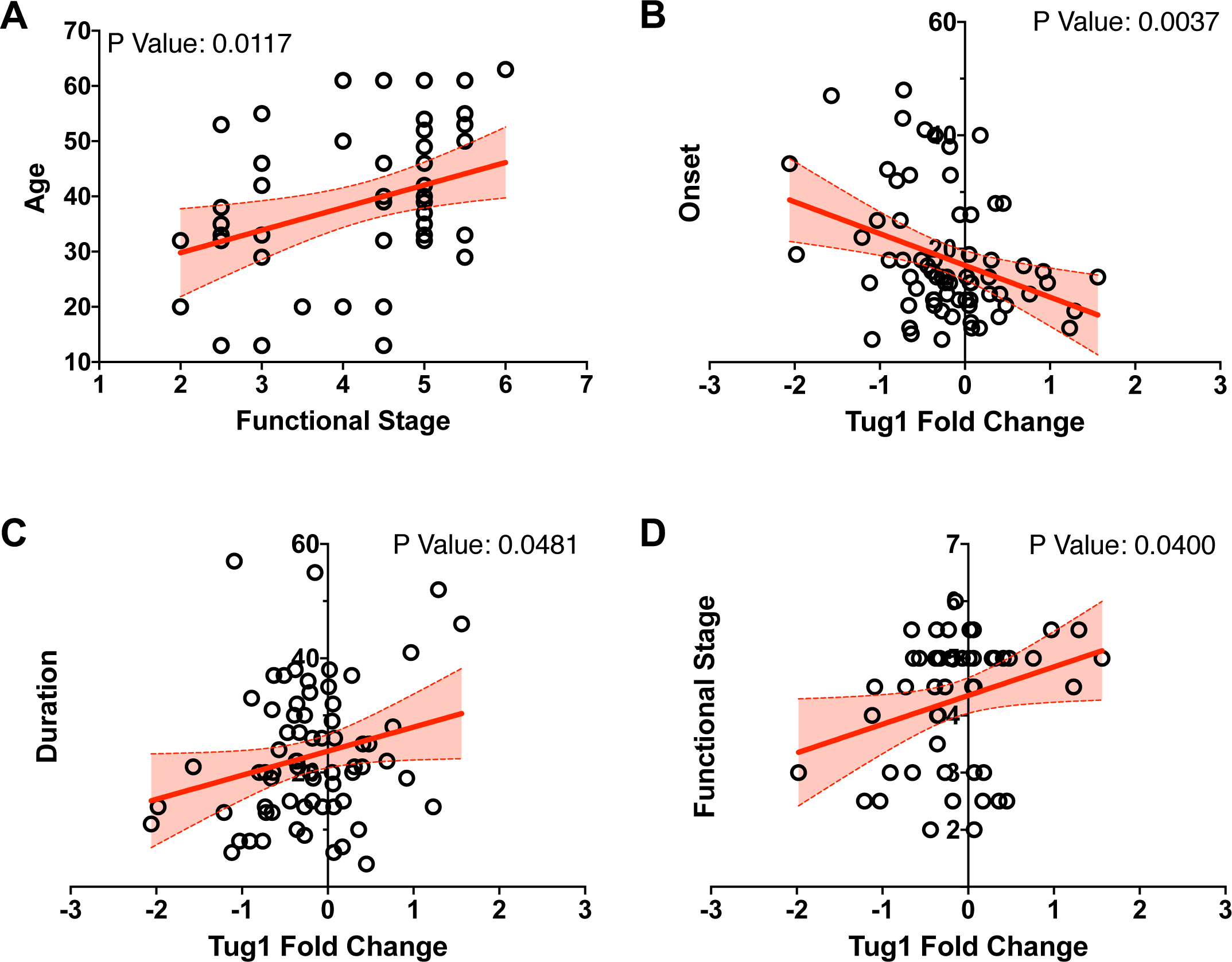
Significant Correlation between TUG1 Expression Levels and Clinical Phenotypes in FRDA Patients. (**A-D**) Scatter plots and linear regression models illustrate the relationship between TUG1 expression levels and various clinical phenotypes. (**A**) As anticipated, a significant association is observed between the functional disability stage and age. (**B**) A significant negative correlation is present between disease onset and TUG1 expression levels (P<0.05). (**C**) A substantial positive correlation exists between TUG1 levels and disease duration (P<0.05). (**D**) The functional disability stage also shows a significant positive correlation with TUG1 levels, highlighting its potential as a biomarker for disease severity.

To further examine the impact of other variables, namely Age, Sex, and GAA expansion (genetic marker of disease severity), on the correlation between TUG1 levels, disease onset, and FDS score, we conducted a multivariate regression analysis. The FDS emerged with an estimated coefficient of 0.1721, hinting at a positive relationship with TUG1 fold change. This infers that an increase in the FDS score is paralleled by a comparable increase in TUG1 levels. However, this association did not reach the threshold of statistical significance (p = 0.0751), calling for further investigation. In contrast, the variable ‘disease onset’ was characterized by an estimated coefficient of -0.02661, representing a statistically significant negative relationship with TUG1 fold change (p = 0.0047). This compelling observation implies that elevated TUG1 levels are associated with earlier disease onset and hence, more genetically severe cases. Residual analysis further validated our regression model, with residuals showing no clear patterns of heteroscedasticity or non-linearity. This supports the robustness of our findings and underscores the significant association between TUG1 levels and disease onset (**Supplementary Figure 3**). Taken together, our findings suggest that patients with increased levels of TUG1 typically experience an earlier disease onset and present with higher FDS scores. On the other hand, patients with lower TUG1 expression tend to have a delayed disease onset and exhibit less severe disease manifestations. These results underscore the potential of TUG1 as a crucial biomarker in the prognosis and management of FRDA.

## Discussion

FRDA is a neurodegenerative disorder characterized by complex molecular mechanisms and limited therapeutic interventions^42^. A significant challenge in the management of FRDA is the rapid monitoring of disease progression and the expedited evaluation of the efficacy of potential treatments^8^. This challenge motivated our study to explore potential blood-based molecular biomarkers for FRDA. Upon re-examining a comprehensive dataset comprising 733 individuals, we identified 293 genes showing differential expression in FRDA patients compared to controls. These genes predominantly function in immune system activities, aligning with existing literature that identifies immune activation as an early pathway regulated following frataxin (Fxn) knockdown^11,12^. This finding not only lends credence to previous studies but also opens avenues for future research in identifying specific biomarkers for FRDA.

Extending the analysis to FRDA knockdown mice, we identified Tug1 and Slc40a1 as particularly promising biomarkers. These genes demonstrated consistent differential expression in both human patients and the FRDAkd mouse model. We validated the expression of Tug1 and Slc40a1 in both the mouse model and human blood samples, underscoring their potential as biomarkers and highlighting the benefits of minimally invasive sample collection. In FRDAkd mice, these genes were validated in different tissues primarily affected in FRDA, such as the heart, dorsal root ganglia neurons, and the cerebellum, among others. Interestingly, Tug1 and Slc40a1 expression was significantly altered as early as two weeks following Fxn knockdown. This suggests the prospective value of these genes as early-stage biomarkers, whose expression is directly influenced by Fxn levels.

The FRDAkd mouse model was instrumental in assessing the correlation between frataxin levels and candidate biomarkers. Importantly, Slc40a1 (ferroportin), involved in iron metabolism linked to FRDA pathology, showed tissue-specific expression changes after Fxn knockdown and partial restoration upon Fxn rescue. This observation correlates with previous findings that have implicated iron metabolism dysregulation in FRDA pathology^43,44^, adding an extra layer of complexity and importance to our study. These collective findings contribute to a deeper understanding of the molecular mechanisms underpinning FRDA and underscore the potential utility of Slc40a1 as therapeutic marker.

Long non-coding RNAs such as Tug1 have multifaceted roles in cellular biology, particularly in the epigenetic regulation of transcription^45–47^. Our findings establish Tug1 as a key molecular player that is downregulated in multiple tissues affected by FRDA pathology, specifically in response to Fxn knockdown. Given that Tug1’s involvement extends to cellular processes like cell proliferation^21–23^, apoptosis^24–28^, and mitochondrial bioenergetics^31^ —processes that are also disrupted in FRDA—the importance of Tug1 in FRDA pathogenesis becomes increasingly evident. We observed a stark downregulation of Tug1 in various tissues, with an exception in liver, upon Fxn knockdown. Strikingly, the levels of Tug1 were partially restored following Fxn recovery, particularly in heart and muscle tissues. This emphasizes Tug1’s potential as a highly specific biomarker for FRDA.

Our analysis further extends to the exploration of Tug1 and Slc40a1 expression in whole blood across eleven time points during both Fxn depletion and recovery stages. Remarkably, Tug1’s strong linear correlation with Fxn levels, especially in comparison to Slc40a1, makes it a more suitable candidate as a peripheral biomarker. The trend of Tug1 expression was statistically significant, showing a linear decrease during the Fxn depletion phase and a linear increase during the Fxn rescue phase. It’s important to note that our findings rule out the confounding influence of dox treatment in FRDAkd mice, strengthening Tug1’s candidacy as a FRDA-specific biomarker. Our study also sheds light on the specific down-regulation of Tug1 in cerebellum, which is crucial given the extensive cerebellar damage observed in FRDA^32,33^. The early detection of Tug1 downregulation in cerebellum and its strong linear correlation with Fxn levels provides compelling evidence for Tug1’s importance in understanding the regional specificity of neuronal damage in FRDA^48–51^. Ultimately, our work delves into the downstream target genes of Tug1, which include genes implicated in cellular processes disrupted in FRDA. For instance, Nedd1, a gene involved in cellular senescence^35^, exhibited significant downregulation in multiple tissues in FRDAkd mice, reinforcing the vital role Tug1 may play in the cellular biology disrupted in FRDA. Another notable gene, Ccnd2, which is implicated in cell cycle regulation^52^, was also known to be regulated by Tug1 and showed tissue-specific alterations. These findings solidify the case for Tug1 not just as a biomarker, but as a critical molecular component in understanding the complex mechanisms underlying FRDA pathogenesis.

Our investigation into TUG1 expression in FRDA patients, heterozygous carriers, and healthy controls, using both public microarray datasets^12^ and reverse transcription-quantitative PCR across multiple cohorts and experimental settings, further consolidates its role as a potential biomarker. The results demonstrated a substantial downregulation of TUG1 in both whole blood and serum of FRDA patients. Moreover, the findings derived from linear and multivariate regression analyses were especially informative. Our data points to a marked association between Tug1 expression levels and clinical variables like disease onset, disease duration, and the FDS score. Notably, the multivariate analysis unveiled that elevated TUG1 levels are strongly linked to earlier disease onset. This significant correlation with early disease onset in FRDA patients provide potential avenues for disease monitoring and therapeutic development. Periodic assessments of Tug1 levels, using easily accessible samples like blood and serum, could serve as tools for tracking disease progression and severity. The implications for therapeutic development are profound, potentially transforming drug development processes and guiding targeted interventions. Insights into the molecular pathways of Tug1 could lead to targeted therapies, transforming FRDA drug development.

Our study employs a comprehensive approach and diverse sample analysis, incorporating both human and mouse model data, to substantiate the validity of our findings. The use of robust validation methods and the FRDAkd mouse model^11^ further enhance the reliability of our study, positioning non-coding RNA Tug1 as a promising biomarker for FRDA. Nevertheless, our study has limitations that warrant attention. Despite our efforts to eliminate confounding effects, the potential impact of unrecognized confounding variables on our results cannot be completely ruled out. Therefore, clinical validation through trials is essential for further substantiation. Additionally, our findings require replication in larger cohorts to establish broader validity, and longitudinal studies are crucial for understanding the long-term association of TUG1 with disease progression. Furthermore, the complex interplay between Fxn, Tug1, and other cellular processes necessitates more detailed investigation.

In conclusion, our rigorous study underscores Tug1’s critical role as a prospective blood-based biomarker for FRDA. Utilizing robust methodology and in-depth analyses, we have confirmed the functional importance of Tug1 and its downstream targets in tissues specific to FRDA. Importantly, our data reveals a strong correlation between Tug1 expression and key clinical indicators, such as disease onset and Functional Disability Stage scores, further establishing its clinical significance. Tug1 stands out as an early-stage, blood-based marker, thereby emphasizing its potential for minimally invasive diagnostic applications with substantial clinical and therapeutic implications. As such, Tug1 offers a promising path for both monitoring the disease and guiding therapeutic development, contributing to improved patient care and deeper understanding of this complex neurodegenerative disorder. Future research should aim to validate these findings through larger and more diverse patient studies while also focusing on the translational potential of these scientific insights into effective clinical applications.

## Supplementary material

Supplementary material is provided as a separate PDF file.

---

**Supplementary Figure 1.**
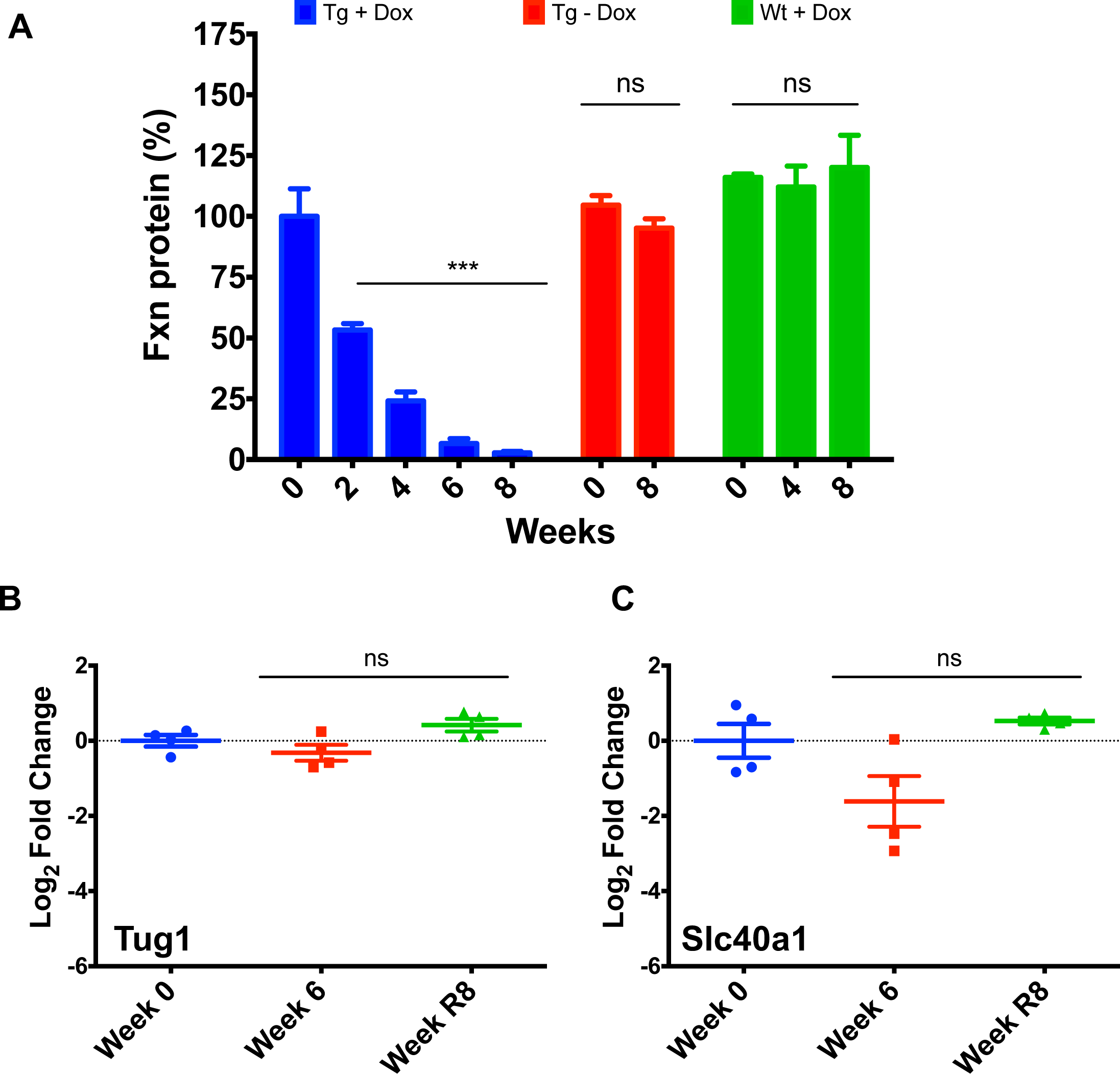
Assessment of FXN Levels in FRDA-Knockdown Mice and Gene Expression Levels in Wildtype Mice. (**A**) Relative quantification of FXN levels in heart tissue samples from FRDA-knockdown mice. Three different groups were tested: Transgenic with doxycycline treatment (Tg + Dox), transgenic without doxycycline treatment (Tg - Dox), and wildtype with doxycycline treatment (Wt + Dox). The FXN levels were measured via ELISA assay. Each group consisted of four samples (N = 4). (**B-C**) Expression levels of (**B**) Tug1 and (**C**) Slc40a1 in wildtype mice treated with doxycycline, followed by a recovery phase. One-way ANOVA and Welch’s t-test were utilized for statistical analyses. Data are presented as mean ± SEM, with *** denoting p ≤ 0.001.

**Supplementary Figure 2.**
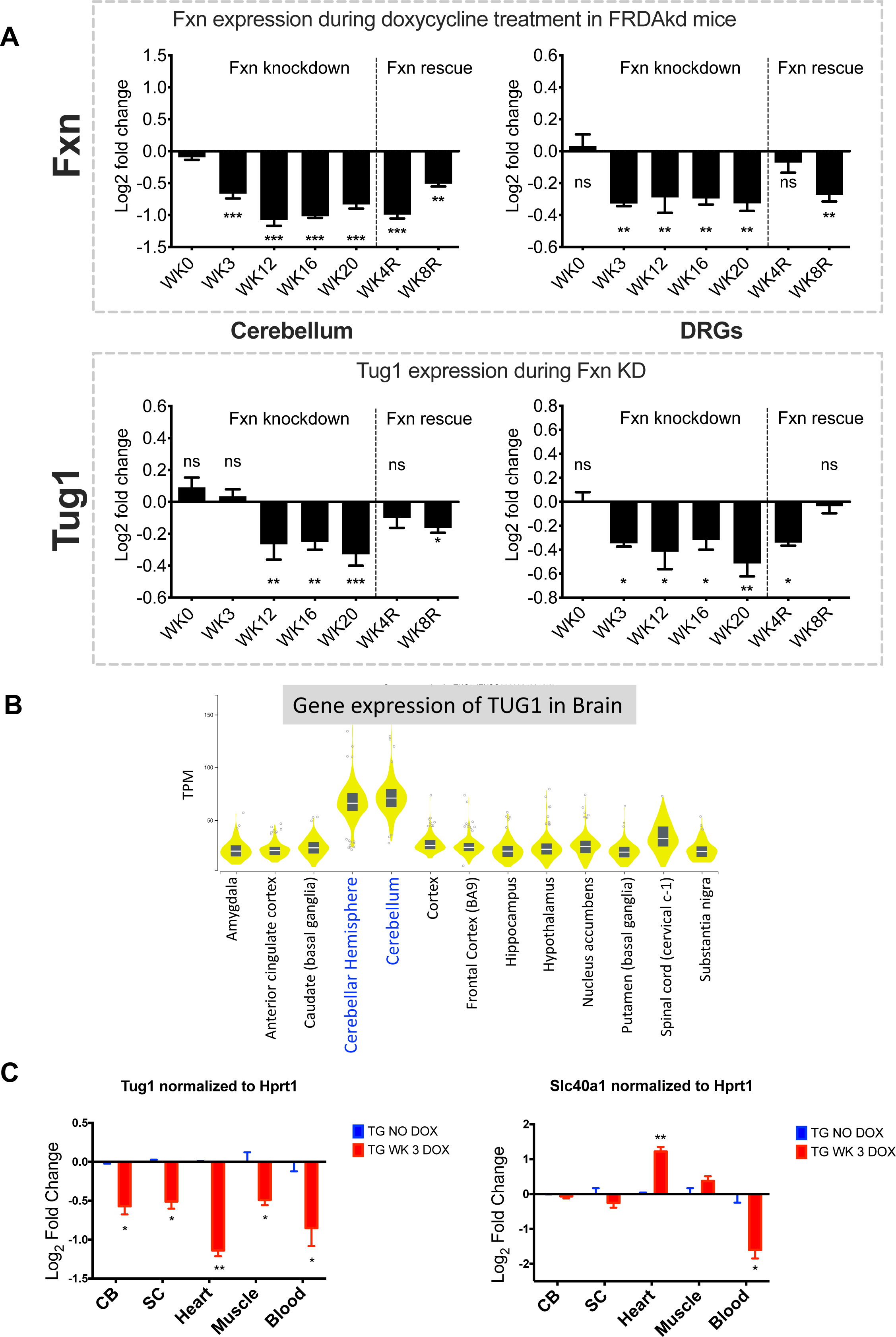
Correlation of Tug1 Expression with Fxn Levels in FRDA-Knockdown Mice and Tug1 Expression across Human Brain Regions. (**A**) Longitudinal analysis of Fxn and Tug1 gene expression in the cerebellum and dorsal root ganglia (DRGs) of FRDA-knockdown mice during frataxin knockdown (at weeks 0, 3, 12, 16, and 20) and recovery (at weeks 4 and 8). Four biological replicates were assessed for each time point. (**B**) Tug1 expression is notably higher in the cerebellum and cerebellar hemispheres, which are critical regions affected in FRDA, as evidenced by data from the GTEx database. The y-axis indicates the normalized expression values in Tags Per Million (TPM), and the x-axis represents various brain regions. (**C**) Expression levels of Tug1 and Slc40a1 in cerebellum, spinal cord, heart, muscle, and blood samples from FRDA-knockdown mice, treated with doxycycline for three weeks, normalized to Hprt1. Three biological replicates (N = 3) were analyzed. Statistical analysis was carried out using Welch’s t-test. Data are presented as mean ± SEM; asterisks denote significance levels as * = p ≤ 0.05, ** = p ≤ 0.01, *** = p ≤ 0.001.

**Supplementary Figure 3:**
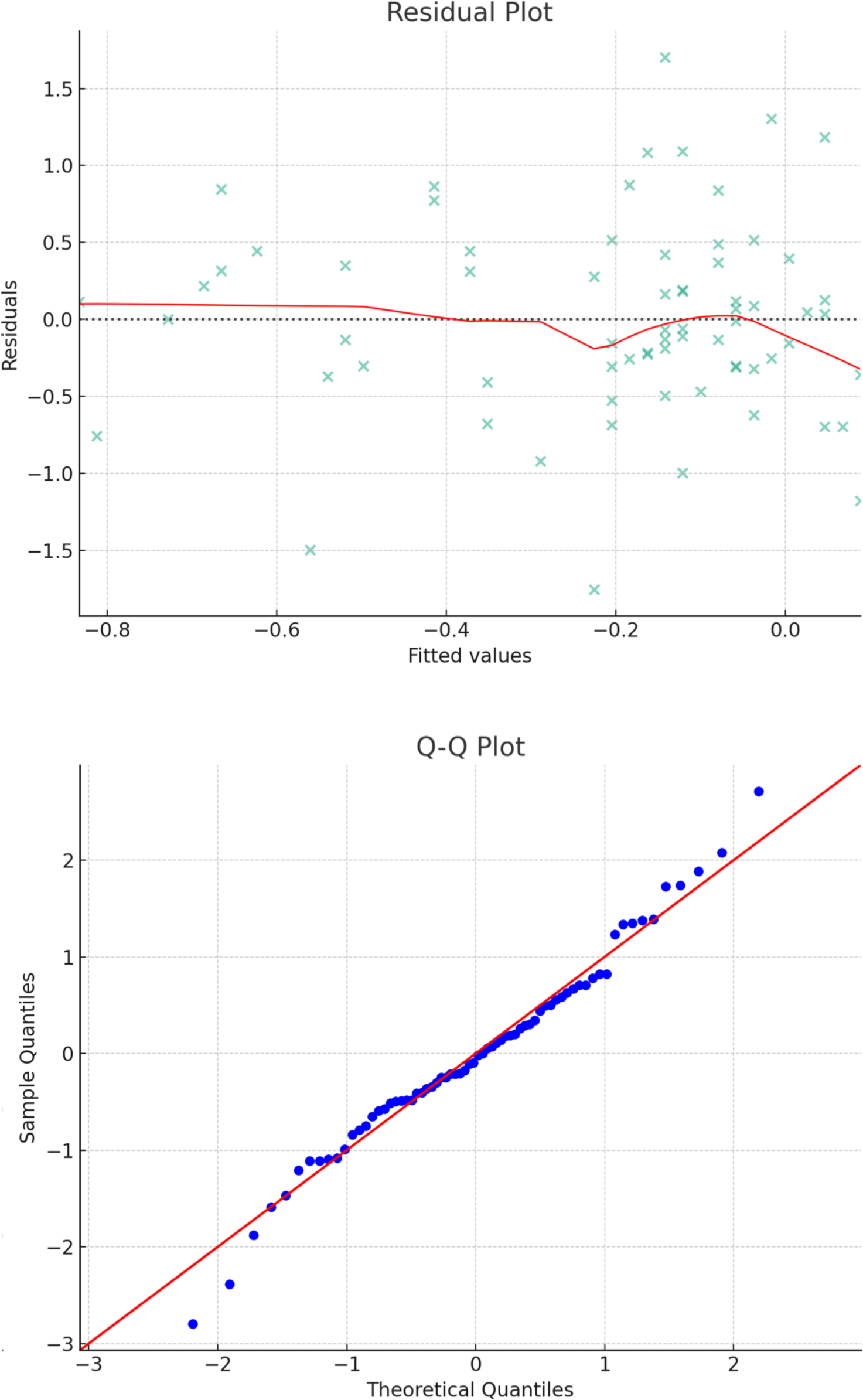
Residual and Normal Q-Q Analysis of Regression Model with TUG1 Levels as the Dependent Variable and Disease Onset as the Independent Variable. The top panel presents a Residual plot, where each data point corresponds to a specific observation plotted against its predicted value. The red Lowess line, ideally lying horizontally at zero if the model assumptions are upheld, exhibits minor deviation, demonstrating an acceptable model fit with no distinct residual patterns. The bottom panel exhibits a Normal Q-Q plot, which juxtaposes the distribution of residuals against the standard normal distribution. The scatter points signify residuals, largely aligning along the diagonal red reference line, thereby implying a predominantly normal distribution of residuals.

